# 3D-Printing in Radiation Oncology: Development and Validation of Custom 3D-Printed Brachytherapy Alignment Device and Phantom

**DOI:** 10.1101/2022.07.03.498548

**Authors:** Destie Provenzano, Hamid Aghdam, Sharad Goyal, Murray Loew, Yuan James Rao

## Abstract

Brachytherapy seeks to treat cancer through insertion of radioactive sources aligned by needles within standardized blocking templates. However, many patients have anatomy that does not conform to standard tools. We created a 3D printed device to provide customized needle alignment for a patient. Device was validated through CT scan and treatment plan creation on a custom anatomical phantom based on patient data. CT scan of a cervix tumor from anonymized patient data was used to develop a 3D printed brachytherapy alignment tool and phantom anatomical mold. Multiple materials were evaluated to match patient anatomy in density and Hounsfield Units present on CT scan, with additional considerations for toxicity, compliance, and practicality. Alignment device and molds were developed in PLA. Silicone of T20 hardness was used to create relevant anatomical organs (Uterus, Rectum, Bladder). Tumor tissue was mimicked by addition of 1CC of Iodine contrast agent to silicone. Device and needles were arranged, inserted into anatomical phantom, and scanned by CT to mimic brachytherapy procedure. 3D printed Silicone uterus of 1.08 g/cm^3 and 40 HU mimicked human uterus on CT scan. Constructed uterus dimensions of 6.5 cm x 5.5 cm x 3.3 cm were verified on imaging to be within + 1 mm of original patient scan. The 1 CC of contrast agent provided sufficient differentiation of “tumor” ring from “tissue.” CT scan and treatment plan creation verified that the alignment device provided correction insertion of needles into the phantom tumor tissue and uterus. This pilot study provides a potential methodology to develop future anatomical phantoms and alignment devices from CT scans of patient data. Additional modifications could make this a viable training tool for future residents and medical students to learn brachytherapy.

**Key Points:**

1. 3D printing can be leveraged to create a Brachytherapy alignment device from patient’s anatomy.
2. Anatomical phantom can be generated from 3D printing for use testing device or for training.
3. Relevant materials to create phantom that mimic patient’s anatomy on CT scan.
4. Generation of a treatment plan based on CT scan was able to validate device and phantom

## I. Introduction

Cervical cancer is the third most common cancer diagnosed in women worldwide, and the ACS predicts there will be 14,480 new cases in the US this year.[1] Brachytherapy is the only method that can deliver the high dose (>80 Gy) required to control Cervical Cancer without side effects. Brachytherapy is a form of radiation therapy that seeks to treat cancer through insertion of radioactive sources at a tumor site. Brachytherapy for Cervical Cancer is typically delivered through interstitial, intracavitary, or combination means that utilize a blocking needle template tool to align interstitial needles to the tumor. Radioactive sources are aligned into the patient by needles and tools such as a blocking needle template tool while undergoing a CT scan to ensure correct placement. However many patients have anatomy that does not conform to standardized existing tools.

3D printing is a manufacturing process that prints a thermoplastic material layer by layer according to a designated X,Y, and Z matrix to create a 3D rendition of an object. Common materials utilized in 3D printing are PLA or PETG, and vary in terms of hardness and heat properties and can be further varied with the actual print through variations in fill density, heat, or other properties. Prior studies have explored the use of 3D printing in Brachytherapy (Table 1). Kadoya et al evaluated the deformation in image registration between external beam radiation and HDR brachytherapy through a 3D printed deformable phantom.[2] The authors created organs by developing a uterus in urethane and creating a 2 mm bladder wall with silicone that was filled with water. Cunningham et al. created an “end to end” phantom using Urethane for the uterus and plaster for bones.[3] They found that acceptable contrast was achievable in CT-SIM and MR-Linac images, However the bladder was challenging to distinguish from the background in CT-SIM. An additional study evaluated the use of a 3D printed head and neck phantom for quality assurance in intensity-modulated radiation therapy (IMRT).[4] All of these models were built for scanning purposes, to date none have tested Brachytherapy inserts.

**Table 1:**
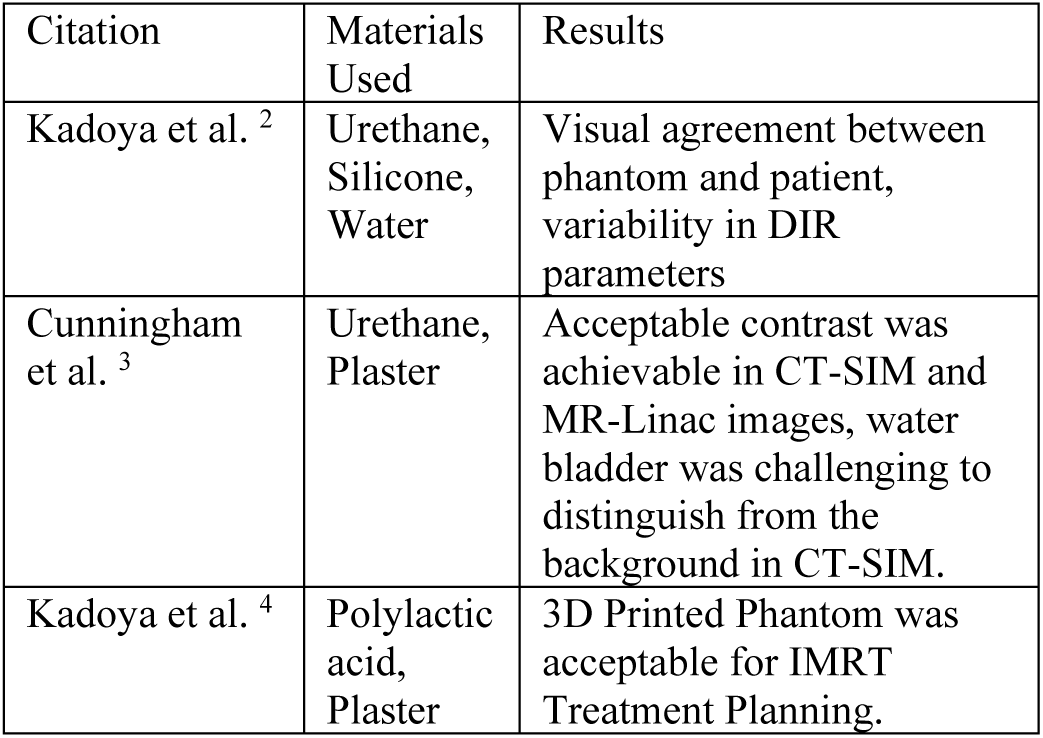
Summary of prior phantoms developed for radiation oncology and brachytherapy purposes.

| Citation | Materials Used | Results |
| --- | --- | --- |
| Kadoya et al. <sup>2</sup> | Urethane, Silicone, Water | Visual agreement between phantom and patient, variability in DIR parameters |
| Cunningham et al. <sup>3</sup> | Urethane, Plaster | Acceptable contrast was achievable in CT-SIM and MR-Linac images, water bladder was challenging to distinguish from the background in CT-SIM. |
| Kadoya et al. <sup>4</sup> | Polylactic acid, Plaster | 3D Printed Phantom was acceptable for IMRT Treatment Planning. |

We sought to create a 3D printed device that provided customized needle alignment for a patient and developed a 3D printed mold and anatomical “phantom” model based on patient CT scan to test the device. Testing of devices such as a custom brachytherapy device needs to be confirmed both via scan (such as a CT scan which is used prior to radiation therapy treatment for patient) and treatment plan creation to ensure device functions as designed. Although there is no standard phantom for brachytherapy for cervix cancer, water filled cubes or meat placed together are often used to model a human. For our purposes we sought to create a 3D printed phantom for testing of custom brachytherapy device and created a treatment plan upon this data to validate correct insertion of needles and relevant proportions of patient anatomy. The primary purpose of creating this tool was for education and to produce simulated treatment plans rather than for clinical treatment planning at this stage.

## II. Methods

### A. Overview

Our process generated 3D models of patient anatomy utilizing free DICOM viewing software (3D Slicer), transported them to a format that could be 3D printed (STL file), evaluated materials based on similarity to the human uterus, (PLA, Silicone), printed the models using a Flashforge Adventure 3D Printer, and finally validated the models with a CT scan of the phantom undergoing a Brachytherapy procedure and treatment plan.

All CT Scans were conducted at GW using the following parameters (Table 2):

**Table 2:** Overview of CT Scan parameters for CT Scans taken at GW hospital of 3D printed device and anatomical phantom.

| Parameter | Description |
| --- | --- |
| CT Scanner Model and Type | GE Optima CT 580 |
| CT Slice Thickness (H&N) | 1.25 mm |
| CT Slice Thickness (SRS Brain) | 1.25 mm |
| H&N | Helical Acquisition |
| SRS Brain | Axial Acquisition |
| Matrix Size | 512x512 |

### B. Digital Model Development

A CT scan of a Cervix tumor from anonymized patient data was used to develop a 3D printed brachytherapy alignment tool and phantom anatomical mold. CT scan was visualized and converted to 3D format within 3D Slicer (4.1.0) software using default parameters for window and leveling. [5,6] Uterus, rectum, and bladder were isolated from patient image and converted to printable STL format using Meshlab (2021.07) software. [7]

After consultation with board certified radiation oncologist, 3D Printed Brachytherapy alignment tool was developed with specific channels to support a combination cervical cancer Brachytherapy treatment into anonymized patient data. 3D Printed Brachytherapy Alignment Tool was developed based on a 30 mm vaginal cylinder with modifications to allow for placement of 9 interstitial needles through the dome of the cylinder. A center channel was included to be used for placement of tandem or interstitial needle. Side channels were included to support placement of interstitial needles into cervix for hybrid intracavitary/ interstitial Brachytherapy approach. Cylinder was selected over ovoids and rings due to the simplicity of this initial design.

A “donut” shaped tumor ring was developed to fit around the base of the uterus and cervix to mimic a cervix tumor and provide a target for insertion of needles.

### C. Material Evaluation

Multiple materials were evaluated to match patient anatomy in density and Hounsfield Units present on CT scan, with additional considerations for toxicity, compliance, and practicality to create the alignment device. Urethane and Silicone were selected as final materials to test the ability to mimic organs due to their use in prior literature for creation of brachytherapy devices and pliability. The human uterus does not have established published density or specific gravity information. As such a pig uterus density (1.06 + 0.01 g/cm3) was used to identify materials for modeling of human uterus. [8] Hounsfield Units of 40.8 HU were used as a target HU for contrast on the CT scan after examination of patient anatomy and established human uterus HU.

Silicone of T20 hardness was selected for organ creation after reviewing and testing multiple materials due to its similarity to pig uterus specific gravity and appearance on CT scan. (Table 3) Polylactic Acid (PLA) printed with a fill density of 85%, where fill density means a measurement of the amount of plastic used on the inside of the print, mimicked 30 HU on CT scan and was used as one potential “holder” for organs to mimic the remainder of the body on the scan. PLA printed with standard 3D printer fill density of 15% was used as an additional “body” mimic to hold the organs and appear different on CT scan. PLA printed with the standard fill density (15%) for 3D printer settings was used to created 3D printed alignment device. The tumor “donut shaped” cylindrical insert was mixed with 1 CC of contrast agent (Gastrografin – Iodine) to get a 1:10 concentration of Contrast to Silicone. This was done to vary the final HU on CT scan.

**Table 3:**
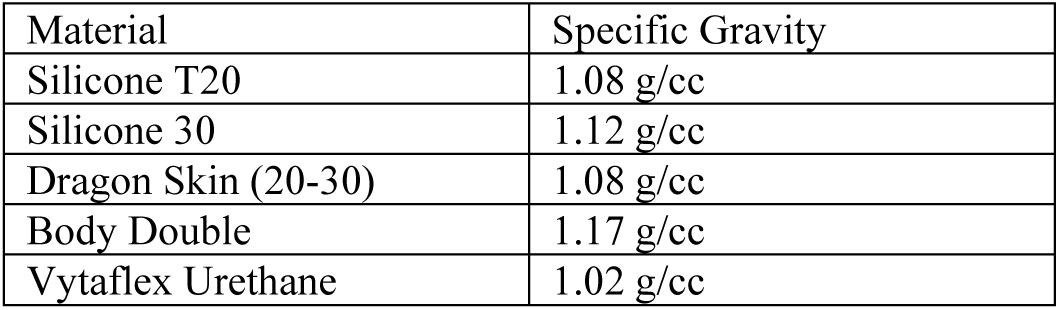
Summary of specific gravity information for relevant materials tested for development of phantom human uterus, bladder, and rectum.

### D. Model Creation

Alignment device and molds were developed and printed in PLA, with silicone of T20 hardness used to create relevant anatomical organs from patient scan (Uterus, Rectum, Bladder) in mold. (Figure 1) Tumor tissue was mimicked by addition of 1CC of Iodine contrast agent to silicone before tumor shape had been molded and set.

**Figure 1:**
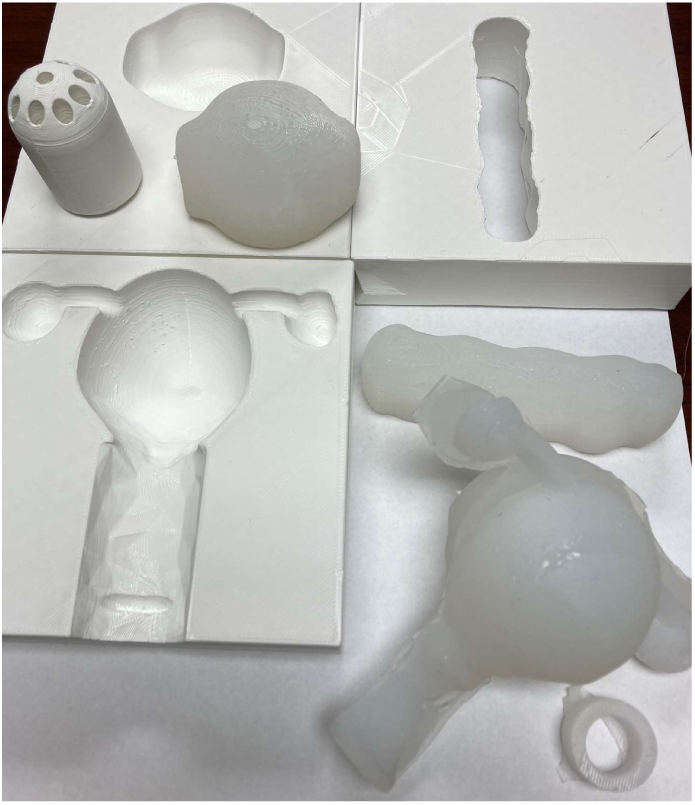
Components of full Brachytherapy Alignment Model. Depiction of “body mimic” containers for organs, silicone organs, Gastrografin contrast agent added to tumor “donut” ring, and 3D printed Brachytherapy alignment device.

**Figure 2:**
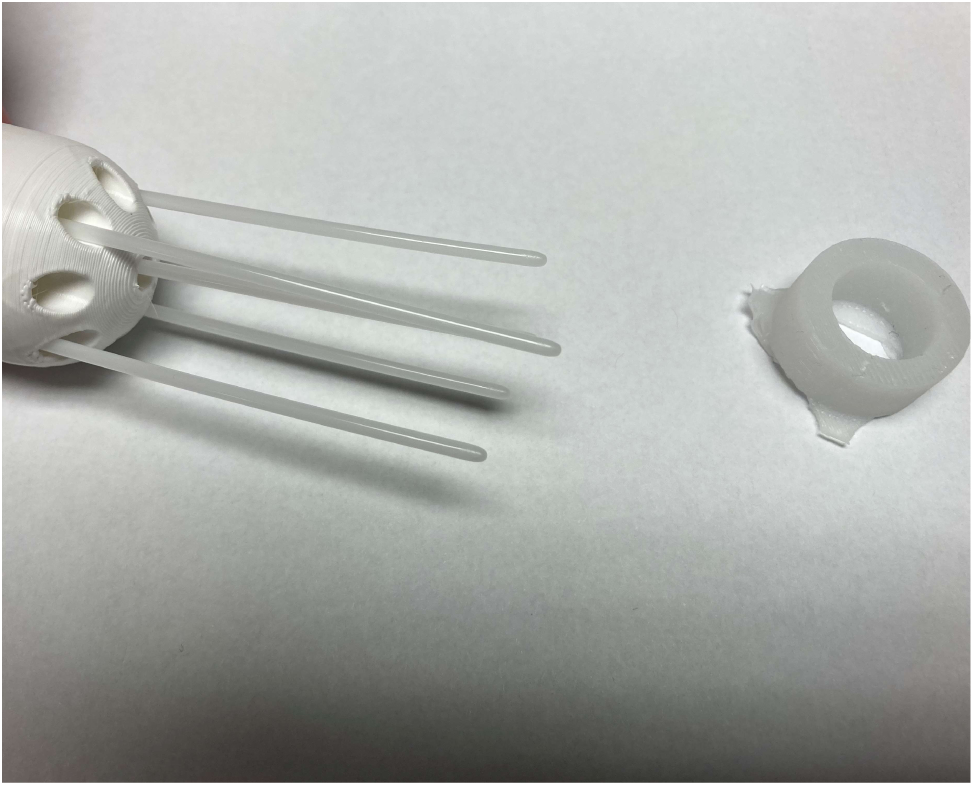
Sample set up of Brachytherapy channels inserted through alignment device.

### E. Validation

Phantom was first arranged and scanned by CT (Figure 3). Device and needles were arranged, inserted into the anatomical phantom, and scanned by CT to mimic the brachytherapy procedure (Figure 4). All needles were set by board certified radiation oncologist and CT scan was run by physicist to mimic the actual procedure. Brachytherapy Treatment Plan was generated by physicist and board certified radiation oncologist to validate device and phantom.

**Figure 3:**
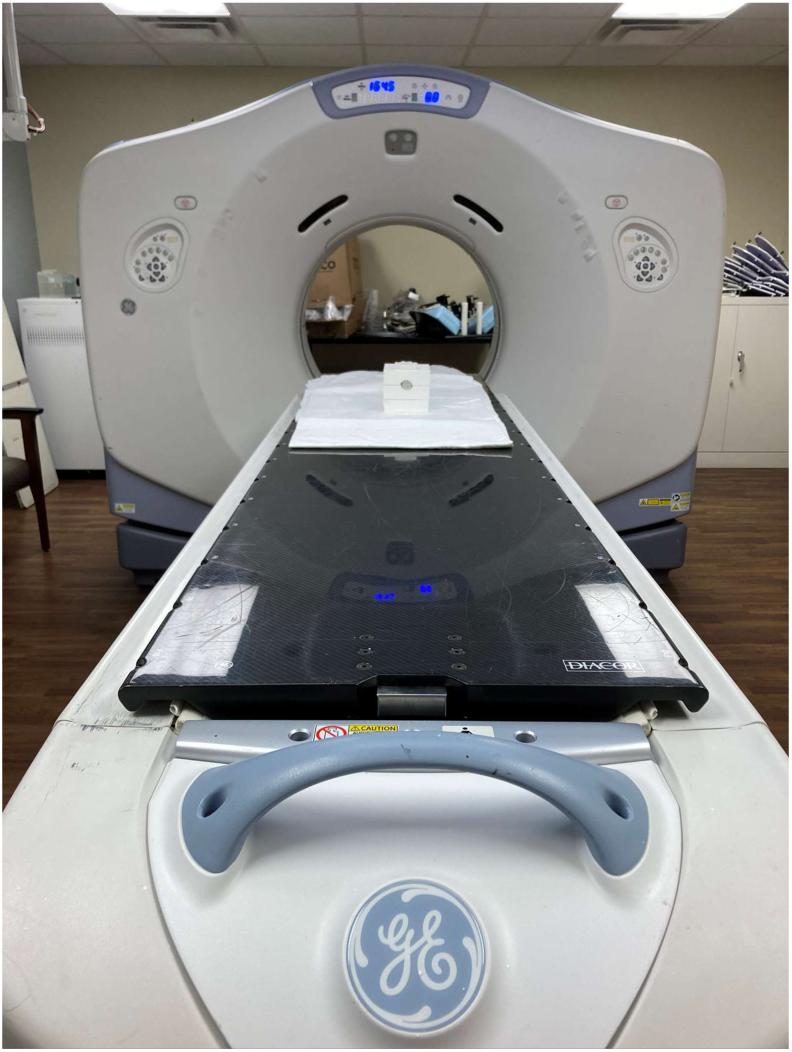
Phantom Aligned on CT bed.

**Figure 4:**
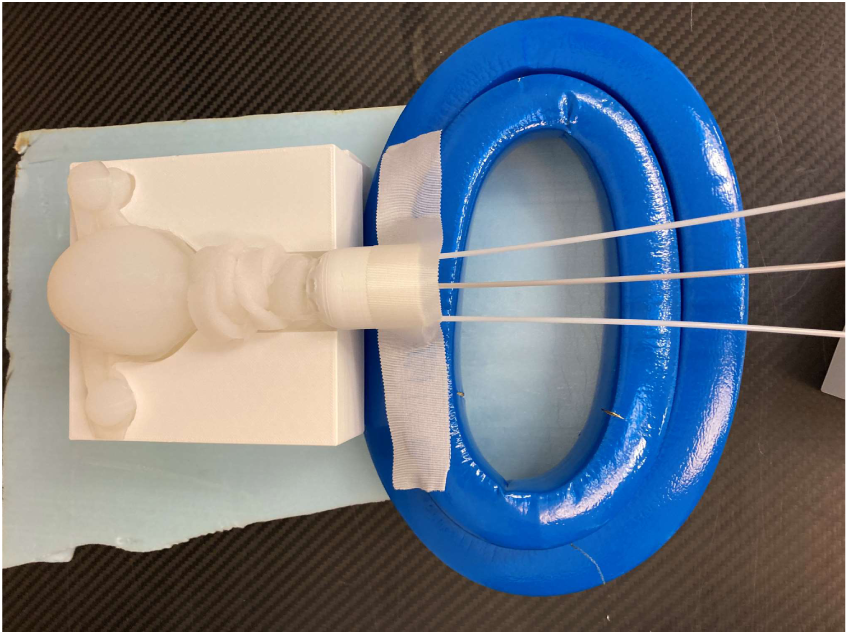
Lower half of assembled 3D Printed silicone model with tumor on cervix and channels inserted into Brachytherapy alignment tool.

**Figure 5:**
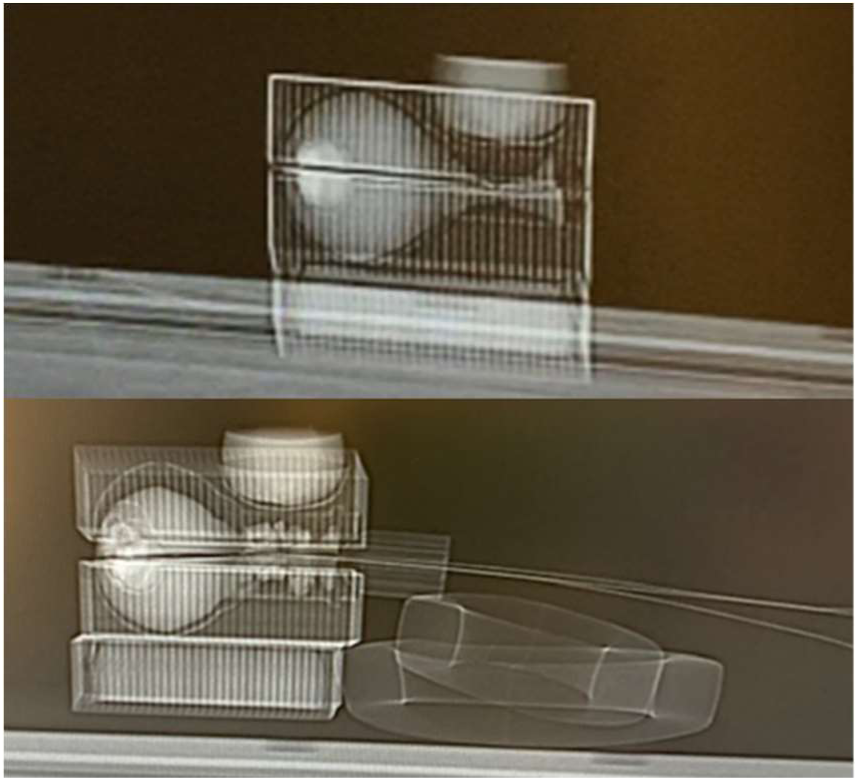
CT Scan of 3d Printed Silicone device with (top) and without (bottom) tumor “donut” placed around cervix area of silicone model. Channels were inserted into tumor “donut” successfully with brachytherapy alignment device guidance (lower).

**Figure 6:**
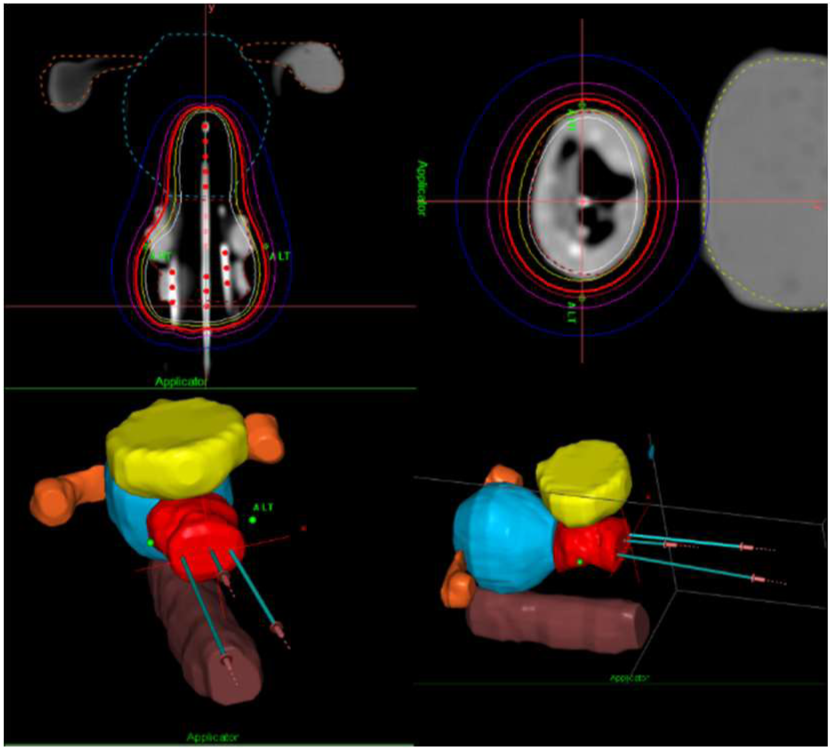
Axial and Sagittal images of treatment plan (Top) and aligned organs and channels (Bottom).

## III. Results

### A. Dimensions of Anatomical Phantom

3D printed Silicone uterus of 1.08 g/cm^3 and 40 HU mimicked human uterus on CT scan, with rectum and bladder also similar to normal in appearance. Constructed uterus dimensions of 6.5 cm x 5.5 cm x 3.3 cm were verified on imaging to be within + 1 mm of original patient scan.

### 3.2 Alignment device and tumor contrast function

Tumor ring around the cervix with 1 CC of contrast agent glowed brighter than tumor tissue, which mimicked scan data. however indicating more CC’s could provide better contrast in the future. Alignment device was able to provide correct insertion of needles into phantom tumor tissue and uterus.

### 3.3 Treatment Plan

Tandem was placed through the center channel into uterus, two side interstitial needles were placed into cervix tumor. For this study we used a thin plastic channel rather than true tandem.

The 3D printed Silicone uterus of 1.08 g/cm^3 and 40 HU mimicked human uterus on a CT scan. Constructed uterus dimensions of 6.5 cm x 5.5 cm x 3.3 cm were verified on imaging to be within + 1 mm of original patient scan. Alignment device was able to provide correct insertion of interstitial needles and tandem into phantom tumor tissue and uterus.

## IV. Discussion

This study aimed to create a pilot anatomical phantom and 3D printed Brachytherapy alignment device for patient anatomy that does not align to standardized templates and tools. An additional anatomical phantom was created to test the device and allow for creation of a treatment plan. This required two steps for validation: 1) validation of the phantom through test insertion of catheters and CT scan, and 2) validation of the device by treatment plan. In both scenarios the anatomical phantom and Brachytherapy alignment device were found to be acceptable from visual inspection of proportions, physical test from insertion of catheters, and CT scan.

Radiation Oncologist was able to insert the catheters into the anatomical phantom, indicating that this could be a useful training tool for future residents and medical students. This also provides a phantom one can make a sample treatment plan upon after insertion of the catheters. Currently there is not an available training tool on the market for Brachytherapy. The alignment device and phantom fulfill this need.

Limitations of this study are that a 3D mold file must be created from patient anatomy in order to test the device. This can add substantial time from the physician contouring the region, or the modeler creating a smoothed successfully printed device. Additionally the PLA device can not be sterilized and would need to be created with a plastic with a higher melting temperature such as PETG for clinical use. Additionally at this time the soler purpose of this study is for training purposes and not for actual treatment planning.

## V. Conclusion

This pilot study provides a potential methodology to develop future anatomical phantoms and alignment devices from CT scans of patient data. Altering concentration of contrast agent could provide better contrast for tumor region. Additional modifications to make models softer could make this a viable training tool for future residents and medical students to learn brachytherapy.

## Abbreviations Used in this Manuscript

CT: Computed Tomography
MRI: Magnetic Resonance Imaging
X-ray: Radiographic Image

